# MiniVess: A dataset of rodent cerebrovasculature from *in vivo* multiphoton fluorescence microscopy imaging

**DOI:** 10.1101/2022.07.19.500542

**Authors:** Charissa Poon, Petteri Teikari, Muhammad Febrian Rachmadi, Henrik Skibbe, Kullervo Hynynen

**Author notes:** corresponding author: Charissa Poon. these authors contributed equally to this work (Charissa Poon, Petteri Teikari).

## Abstract

We present MiniVess, the first annotated dataset of rodent cerebrovasculature, acquired using two-photon fluorescence microscopy. MiniVess consists of 70 3D image volumes with segmented ground truths. Segmentations were created using traditional image processing operations, a U-Net, and manual proofreading. Code for image preprocessing steps and the U-Net are provided. Supervised machine learning methods have been widely used for automated image processing of biomedical images. While much emphasis has been placed on the development of new network architectures and loss functions, there has been an increased emphasis on the need for publicly available annotated, or segmented, datasets. Annotated datasets are necessary during model training and validation. In particular, datasets that are collected from different labs are necessary to test the generalizability of models. We hope this dataset will be helpful in testing the reliability of machine learning tools for analyzing biomedical images.

## Background & Summary

Blood vessel segmentation is often a necessary prerequisite for extracting meaningful analyses from biomedical imaging data. By creating a segmentation, or a mask, that separates vascular from non-vascular pixels, structural information about the vascular system can be acquired, such as diameter, branch order, and blood vessel type. Identification of blood vessels as arterioles, venules, or capillaries can be used to analyze vascular dynamics, such as blood flow and vascular supply. Blood vessel segmentation has clear clinical value. For example, in ischemic stroke studies, vascular segmentation enables detection and quantification of vascular occlusions, which can be helpful in determining therapeutic options.^1,2^. Structural characteristics can also be used as predictors or markers to assist in the diagnosis of diseases, such as Alzheimer’s disease^3,4^, traumatic brain injury^5^, brain tumours^6^, atherosclerosis^7^, and retinal pathology^8,9^.

Apart from vascular analyses, blood vessel segmentation is also a necessary preprocessing step for the analysis of cells and pathological entities (Figure 1). In addition to the endothelial and mural cells that make up the blood vessel proper, various other cell types interact with vascular walls, including astrocyte endfeet processes, perivascular macrophages, and peripheral leukocytes. Such cells and their interactions with vasculature can be identified and analyzed based on distance metrics to vascular walls, a task which is simplified with accurate vascular segmentation masks. Vascular-cellular interactions have been of particular interest in studies focused on diseases. For example, recruitment of peripheral leukocytes to cerebrovasculature has been observed following traumatic brain injury^10^, middle cerebral artery occlusion^11^, and in Alzheimer’s disease^12^. Similar distance metrics can be used to analyze pathological entities, such as perivascular Aβ plaques^13^ and atherosclerotic plaques^14^. Thus, segmentation of blood vessels is a necessary preprocessing step that facilitates further vascular and cellular analyses.

**Figure 1.**
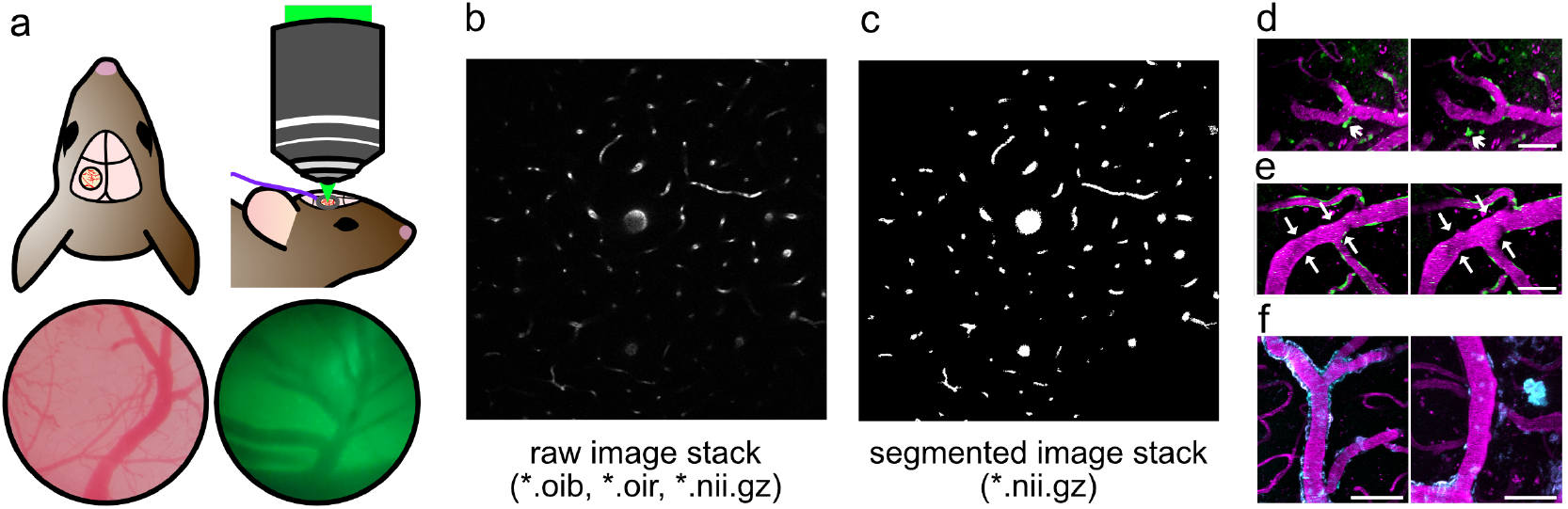
General workflow for 2PFM image processing. a) A cranial window was created on the parietal bone, enabling *in vivo* 2PFM imaging. Superficial blood vessels can be observed through the cranial window (left) and ocular lens of the microscope (right). b) Raw image volumes of rodent cerebrovasculature were collected and saved in Olympus image formats (*.oib, *.oir). c) Segmentations of blood vessels were achieved using a U-Net and manual corrections. In the MiniVess dataset, raw and segmented image volumes are shared as NIfTI files (*.nii.gz). Segmentation of blood vessels enables downstream analyses, such as (d) cell tracking (arrow heads), (e) vasoconstriction and dilation (arrows), and (f) analysis of pathological entities, such as Aβ plaques (cyan). Blood vessels are shown in magenta. Scale bars = 50 μm.

In the neurosciences, two-photon fluorescence microscopy (2PFM) is currently the technique of choice for intravital microscopy. While the resolution of 2PFM can be on par with confocal microscopy, the risk of phototoxicity and photobleaching of tissues and fluorophores is substantially reduced because the excitation volume is limited to the focal volume of the microscope^15^. The use of longer wavelengths also results in less scattering by the neural tissue, allowing imaging at deeper depths within the brain. 2PFM has been extensively used to investigate various phenomena, including neural activity using voltage-sensitive dyes^16^and calcium indicators^17^, microglial activity using transgenic animal models^18^, and vascular dynamics^19^.

Most methods of automated image processing of 2PFM images rely on proprietary software, such as Imaris (Bitplane, United Kingdom) and Volocity (Quorum Technologies, Canada). Each analysis type, such as vascular segmentation and cell tracking, is generally sold as separate modules. Comprehensive analyses of datasets are therefore functionally limited by the modules available and can become prohibitively expensive. Furthermore, while automated modules produce impressive results for images with high SNR, biomedical images, particularly intravital 2PFM images, are inherently noisy. In practice, substantial manual modifications are required. An open-source alternative is FIJI (Fiji Is Just ImageJ)^20^. However, FIJI plugins often lack extensive documentation, resulting in a ’black-box’ nature that may deter and limit use.

Deep learning, such as convolutional neural networks (CNNs) and recent transformer-based model architectures^21,22^, have been extensively used for automated segmentation tasks in biomedical imaging. For example, the U-Net, a fully CNN, achieves impressive performance in segmenting densely packed neurons in electron microscopy images^23^. Clinically, CNNs have achieved state-of-the-art performances in segmenting brain vasculature in magnetic resonance angiography^24^ and retinal vasculature in optical coherence tomography angiography^25^ datasets, which have been used to assist in the identification of pathological features.

A common challenge in the application of deep learning models to the biomedical imaging field is the generalizability of models. Models are often exclusively trained on datasets that were collected from a single site. Such models often fail to perform when evaluated on datasets collected at different sites due to a so-called ’domain shift’ (see e.g. Ouyang *et al*. 2021^26^ for an example in medical image segmentation), caused by differences in tissue preparation, scanner or microscope set-up, and/or inter-user variability in defining labels^27,28^. The problem is compounded by poor reporting of the number of evaluation sites and samples used^29^. One way to improve the reliability and transparency of ML models is to use diverse samples during training, and independent data cohorts for testing^30^. However, the availability of such annotated, publicly available biomedical imaging datasets is limited due to ethical and privacy concerns, particularly in clinical studies. Another strategy is to use synthetic datasets or publicly available non-biomedical datasets (e.g. ImageNet) as part of the training process, and then evaluate the trained model on the real dataset, a process known as ’transfer learning’^31,32^. For example, using transfer learning, a CNN that was pre-trained on a synthetic dataset of blood vessels resulted in impressive segmentation of real mouse brain vasculature^33^. However, the availability of real, annotated, field-specific datasets remains to be a need for evaluating the generalizability of models in the biomedical imaging field. In addition, there has been a recent shift in focus from adjusting model parameters to achieve better performance metrics (’model-centric’), to improving the quality of datasets to improve performance metrics (’data-centric’), highlighting the importance of high-quality, publicly available datasets.

Public microscopy datasets have been curated by various research groups world-wide. For example, the Human Protein Atlas shows the distribution and expression of proteins and genes across major organ systems^34–>36^, the Broad Bioimage Benchmark Collection contains annotated cell datasets^37^, and the Allen Brain Cell Types Atlas offers electrophysiological, morphological, and transcriptomic data measured from human and mouse brain. However, vascular datasets have not been as extensively documented. The availability of an annotated 2PFM vascular dataset would assist in diversifying the samples used for training a segmentation model, or in evaluating the performance of segmentation models that were trained on other datasets.

We hereby present MiniVess, an expert-annotated dataset of 70 3D 2PFM image volumes of rodent cerebrovasculature. The dataset can be used for training segmentation networks^38,39^, fine-tuning self-supervised pre-trained networks^>31,>32,>40^, and as an external validation set for assessing a model’s generalizability^41^. The 3D volumes in this dataset have been curated to only contain clean XYZ imaging in order to ensure correct and consistent annotations, or segmentations, which has been observed to be integral to the evaluation of machine learning models^42^. Code for image preprocessing and the U-Net workflow are also provided in the MiniVess project Github page. The U-Net code was written using MONAI, a PyTorch-based framework that was built to encourage best practices for AI development in healthcare research. We hope that the availability of the image volumes and code will assist in evaluating the reliability of models built for the analysis of biomedical images.

## Methods

### Animal preparation

This dataset consists of 2PFM images of the cortical vasculature in adult male and female mice from the C57BL/6 and CD1 strains (20-30 g), and EGFP Wistar rats (Wistar-TgN(CAG-GFP)184ys) (310-630 g)^43^. All animal procedures were approved and conducted in compliance with the Animal Care Committee guidelines at Sunnybrook Research Institute, Canada.

To allow optical access to the brain, an acute cranial window was created over the parietal bone (Figure 1). Detailed protocols on cranial window procedures have been published elsewhere^44^. Briefly, animals were anesthetized using 1.5-2% isoflurane in a mix of medical air and oxygen. Following fur and scalp removal, a 3-4 mm circle (mice) or rectangle (rats) of bone was removed from the parietal bone using a dental drill, and replaced with a glass cover slip. Due to the thickness of the skull in rats, 1% agarose was deposited onto the brain to prevent air bubbles beneath the cover slip. Animal physiology was monitored using a pulse oximeter, and temperature was maintained using a heating pad with a rectal thermistor. To visualize vasculature, Texas Red 70 kDa dextran (dissolved in PBS, 5 mg/kg; Invitrogen, Canada) was injected through a tail vein catheter. Animals were sacrificed under deep anesthesia using cervical dislocation (mice) or euthanol injection (rats) following the end of imaging.

### Imaging

Imaging was conducted using a FV1000MPE multiphoton laser scanning microscope (Olympus Corp., Japan) with an InSight DS tunable laser (Spectra-Physics, USA), or a Ti:Sa laser (MaiTai, Spectra-Physics, Germany). A 25× water-immersion objective lens (XLPN25XWMP2, NA 1.05, WD 2 mm, Olympus Corp., Japan) was used to collect 512 × 512 images with a lateral resolution of 0.621-0.994 *μ*m/pixel, an imaging speed of 2-8 *μ*s/pixel, and a step-size of 1-10 *μ*m, for a maximum depth of 700 *μ*m. Excitation wavelengths of 810 or 900 nm were used. Fluorescent emissions were collected with photo-multiplier tubes or gallium arsenide phosphide (GaAsP) detectors. Images were saved in Olympus’s .oib or .oir file formats. Image details are listed in Online-only Table 1.

**Table 1.**
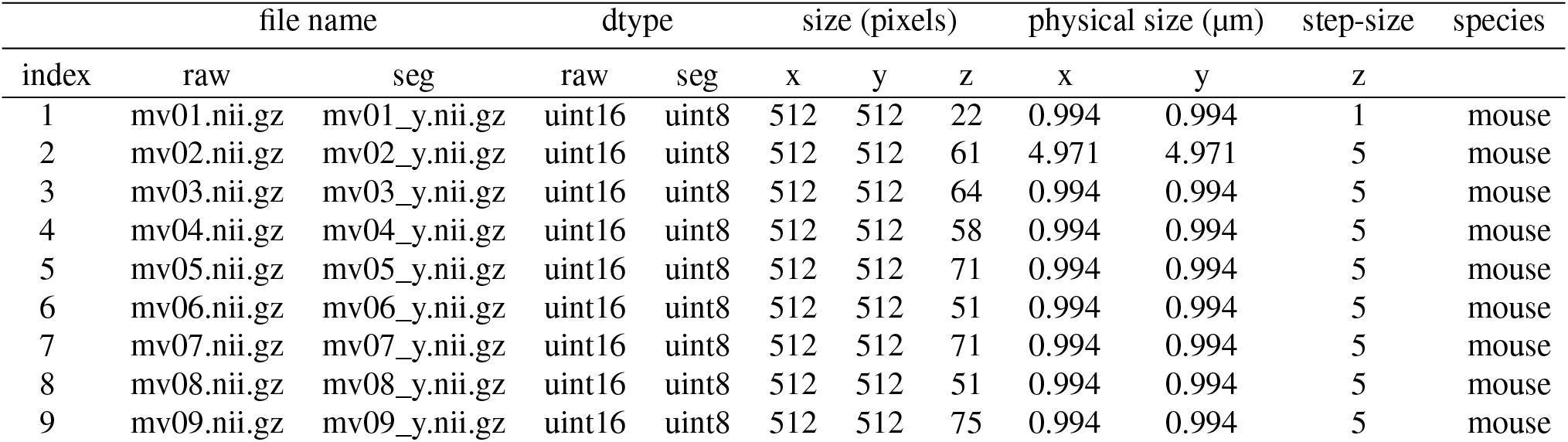
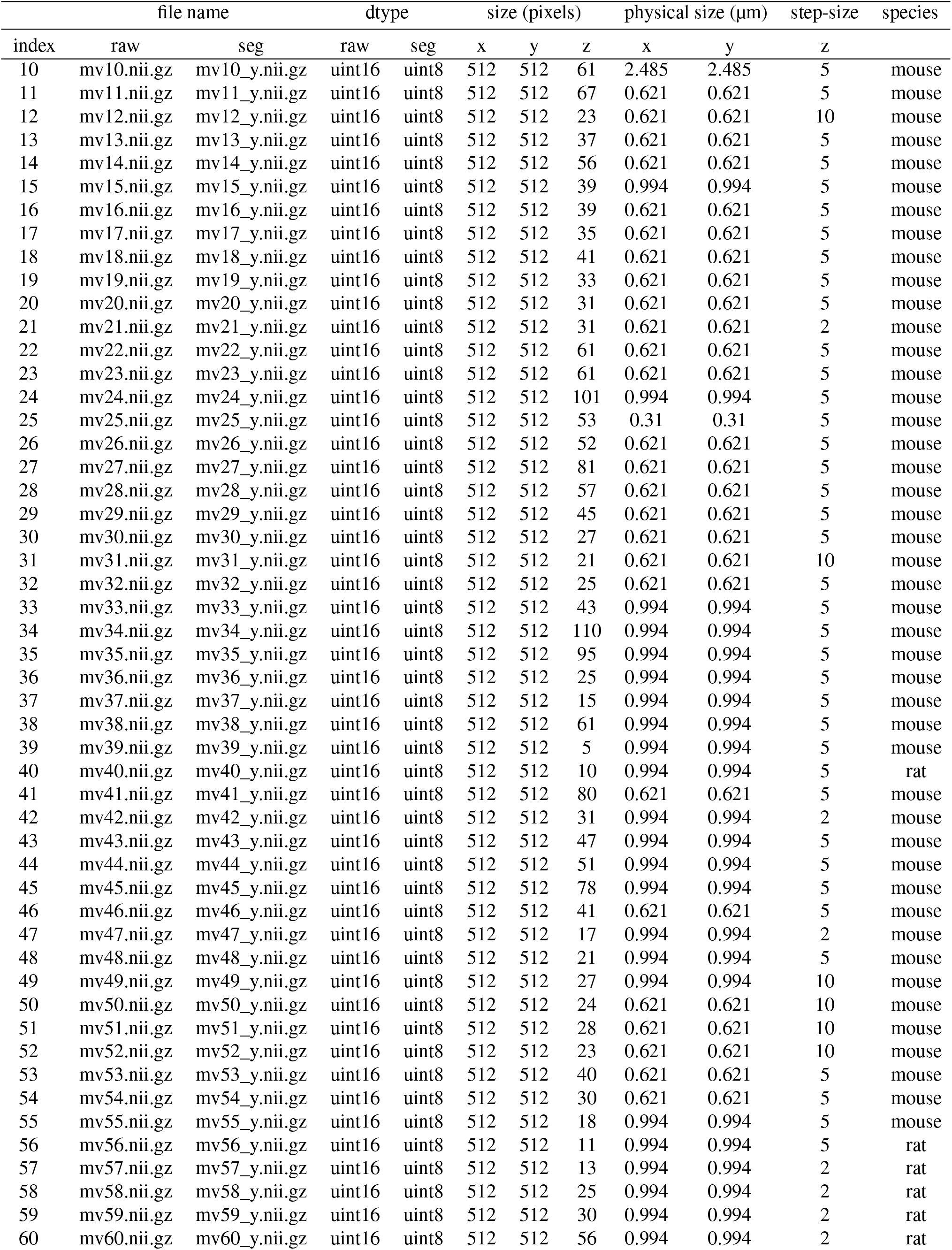
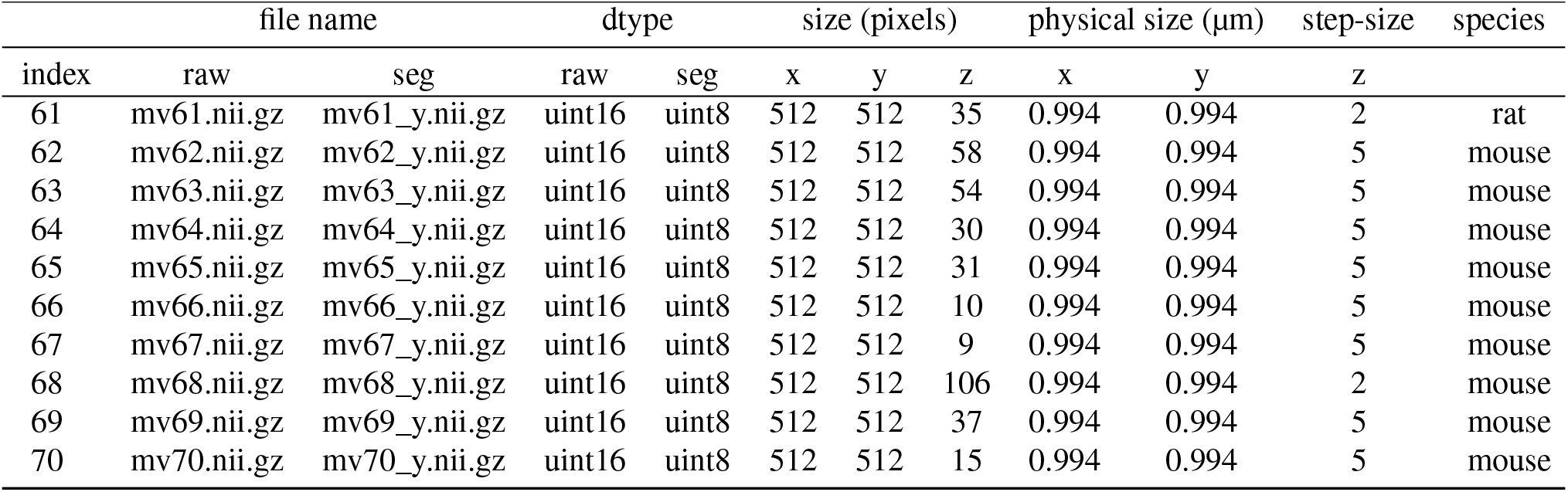
Details of raw and segmented image volumes.

| file name |  |  | dtype |  | size (pixels) |  |  | physical size (μm) |  | step-size | species |
| --- | --- | --- | --- | --- | --- | --- | --- | --- | --- | --- | --- |
| index | raw | seg | raw | seg | x | y | z | x | y | z |  |
| 1 | mv01.nii.gz | mv01_y.nii.gz | uint16 | uint8 | 512 | 512 | 22 | 0.994 | 0.994 | 1 | mouse |
| 2 | mv02.nii.gz | mv02_y.nii.gz | uint16 | uint8 | 512 | 512 | 61 | 4.971 | 4.971 | 5 | mouse |
| 3 | mv03.nii.gz | mv03_y.nii.gz | uint16 | uint8 | 512 | 512 | 64 | 0.994 | 0.994 | 5 | mouse |
| 4 | mv04.nii.gz | mv04_y.nii.gz | uint16 | uint8 | 512 | 512 | 58 | 0.994 | 0.994 | 5 | mouse |
| 5 | mv05.nii.gz | mv05_y.nii.gz | uint16 | uint8 | 512 | 512 | 71 | 0.994 | 0.994 | 5 | mouse |
| 6 | mv06.nii.gz | mv06_y.nii.gz | uint16 | uint8 | 512 | 512 | 51 | 0.994 | 0.994 | 5 | mouse |
| 7 | mv07.nii.gz | mv07_y.nii.gz | uint16 | uint8 | 512 | 512 | 71 | 0.994 | 0.994 | 5 | mouse |
| 8 | mv08.nii.gz | mv08_y.nii.gz | uint16 | uint8 | 512 | 512 | 51 | 0.994 | 0.994 | 5 | mouse |
| 9 | mv09.nii.gz | mv09_y.nii.gz | uint16 | uint8 | 512 | 512 | 75 | 0.994 | 0.994 | 5 | mouse |
| index | raw | seg | raw | seg | x | y | z | x | y | z |  |
| 10 | mv10.nii.gz | mv10_y.nii.gz | uint16 | uint8 | 512 | 512 | 61 | 2.485 | 2.485 | 5 | mouse |
| 11 | mv11.nii.gz | mv11_y.nii.gz | uint16 | uint8 | 512 | 512 | 67 | 0.621 | 0.621 | 5 | mouse |
| 12 | mv12.nii.gz | mv12_y.nii.gz | uint16 | uint8 | 512 | 512 | 23 | 0.621 | 0.621 | 10 | mouse |
| 13 | mv13.nii.gz | mv13_y.nii.gz | uint16 | uint8 | 512 | 512 | 37 | 0.621 | 0.621 | 5 | mouse |
| 14 | mv14.nii.gz | mv14_y.nii.gz | uint16 | uint8 | 512 | 512 | 56 | 0.621 | 0.621 | 5 | mouse |
| 15 | mv15.nii.gz | mv15_y.nii.gz | uint16 | uint8 | 512 | 512 | 39 | 0.994 | 0.994 | 5 | mouse |
| 16 | mv16.nii.gz | mv16_y.nii.gz | uint16 | uint8 | 512 | 512 | 39 | 0.621 | 0.621 | 5 | mouse |
| 17 | mv17.nii.gz | mv17_y.nii.gz | uint16 | uint8 | 512 | 512 | 35 | 0.621 | 0.621 | 5 | mouse |
| 18 | mv18.nii.gz | mv18_y.nii.gz | uint16 | uint8 | 512 | 512 | 41 | 0.621 | 0.621 | 5 | mouse |
| 19 | mv19.nii.gz | mv19_y.nii.gz | uint16 | uint8 | 512 | 512 | 33 | 0.621 | 0.621 | 5 | mouse |
| 20 | mv20.nii.gz | mv20_y.nii.gz | uint16 | uint8 | 512 | 512 | 31 | 0.621 | 0.621 | 5 | mouse |
| 21 | mv21.nii.gz | mv21_y.nii.gz | uint16 | uint8 | 512 | 512 | 31 | 0.621 | 0.621 | 2 | mouse |
| 22 | mv22.nii.gz | mv22_y.nii.gz | uint16 | uint8 | 512 | 512 | 61 | 0.621 | 0.621 | 5 | mouse |
| 23 | mv23.nii.gz | mv23_y.nii.gz | uint16 | uint8 | 512 | 512 | 61 | 0.621 | 0.621 | 5 | mouse |
| 24 | mv24.nii.gz | mv24_y.nii.gz | uint16 | uint8 | 512 | 512 | 101 | 0.994 | 0.994 | 5 | mouse |
| 25 | mv25.nii.gz | mv25_y.nii.gz | uint16 | uint8 | 512 | 512 | 53 | 0.31 | 0.31 | 5 | mouse |
| 26 | mv26.nii.gz | mv26_y.nii.gz | uint16 | uint8 | 512 | 512 | 52 | 0.621 | 0.621 | 5 | mouse |
| 27 | mv27.nii.gz | mv27_y.nii.gz | uint16 | uint8 | 512 | 512 | 81 | 0.621 | 0.621 | 5 | mouse |
| 28 | mv28.nii.gz | mv28_y.nii.gz | uint16 | uint8 | 512 | 512 | 57 | 0.621 | 0.621 | 5 | mouse |
| 29 | mv29.nii.gz | mv29_y.nii.gz | uint16 | uint8 | 512 | 512 | 45 | 0.621 | 0.621 | 5 | mouse |
| 30 | mv30.nii.gz | mv30_y.nii.gz | uint16 | uint8 | 512 | 512 | 27 | 0.621 | 0.621 | 5 | mouse |
| 31 | mv31.nii.gz | mv31_y.nii.gz | uint16 | uint8 | 512 | 512 | 21 | 0.621 | 0.621 | 10 | mouse |
| 32 | mv32.nii.gz | mv32_y.nii.gz | uint16 | uint8 | 512 | 512 | 25 | 0.621 | 0.621 | 5 | mouse |
| 33 | mv33.nii.gz | mv33_y.nii.gz | uint16 | uint8 | 512 | 512 | 43 | 0.994 | 0.994 | 5 | mouse |
| 34 | mv34.nii.gz | mv34_y.nii.gz | uint16 | uint8 | 512 | 512 | 110 | 0.994 | 0.994 | 5 | mouse |
| 35 | mv35.nii.gz | mv35_y.nii.gz | uint16 | uint8 | 512 | 512 | 95 | 0.994 | 0.994 | 5 | mouse |
| 36 | mv36.nii.gz | mv36_y.nii.gz | uint16 | uint8 | 512 | 512 | 25 | 0.994 | 0.994 | 5 | mouse |
| 37 | mv37.nii.gz | mv37_y.nii.gz | uint16 | uint8 | 512 | 512 | 15 | 0.994 | 0.994 | 5 | mouse |
| 38 | mv38.nii.gz | mv38_y.nii.gz | uint16 | uint8 | 512 | 512 | 61 | 0.994 | 0.994 | 5 | mouse |
| 39 | mv39.nii.gz | mv39_y.nii.gz | uint16 | uint8 | 512 | 512 | 5 | 0.994 | 0.994 | 5 | mouse |
| 40 | mv40.nii.gz | mv40_y.nii.gz | uint16 | uint8 | 512 | 512 | 10 | 0.994 | 0.994 | 5 | rat |
| 41 | mv41.nii.gz | mv41_y.nii.gz | uint16 | uint8 | 512 | 512 | 80 | 0.621 | 0.621 | 5 | mouse |
| 42 | mv42.nii.gz | mv42_y.nii.gz | uint16 | uint8 | 512 | 512 | 31 | 0.994 | 0.994 | 2 | mouse |
| 43 | mv43.nii.gz | mv43_y.nii.gz | uint16 | uint8 | 512 | 512 | 47 | 0.994 | 0.994 | 5 | mouse |
| 44 | mv44.nii.gz | mv44_y.nii.gz | uint16 | uint8 | 512 | 512 | 51 | 0.994 | 0.994 | 5 | mouse |
| 45 | mv45.nii.gz | mv45_y.nii.gz | uint16 | uint8 | 512 | 512 | 78 | 0.994 | 0.994 | 5 | mouse |
| 46 | mv46.nii.gz | mv46_y.nii.gz | uint16 | uint8 | 512 | 512 | 41 | 0.621 | 0.621 | 5 | mouse |
| 47 | mv47.nii.gz | mv47_y.nii.gz | uint16 | uint8 | 512 | 512 | 17 | 0.994 | 0.994 | 2 | mouse |
| 48 | mv48.nii.gz | mv48_y.nii.gz | uint16 | uint8 | 512 | 512 | 21 | 0.994 | 0.994 | 5 | mouse |
| 49 | mv49.nii.gz | mv49_y.nii.gz | uint16 | uint8 | 512 | 512 | 27 | 0.994 | 0.994 | 10 | mouse |
| 50 | mv50.nii.gz | mv50_y.nii.gz | uint16 | uint8 | 512 | 512 | 24 | 0.621 | 0.621 | 10 | mouse |
| 51 | mv51.nii.gz | mv51_y.nii.gz | uint16 | uint8 | 512 | 512 | 28 | 0.621 | 0.621 | 10 | mouse |
| 52 | mv52.nii.gz | mv52_y.nii.gz | uint16 | uint8 | 512 | 512 | 23 | 0.621 | 0.621 | 10 | mouse |
| 53 | mv53.nii.gz | mv53_y.nii.gz | uint16 | uint8 | 512 | 512 | 40 | 0.621 | 0.621 | 5 | mouse |
| 54 | mv54.nii.gz | mv54_y.nii.gz | uint16 | uint8 | 512 | 512 | 30 | 0.621 | 0.621 | 5 | mouse |
| 55 | mv55.nii.gz | mv55_y.nii.gz | uint16 | uint8 | 512 | 512 | 18 | 0.994 | 0.994 | 5 | mouse |
| 56 | mv56.nii.gz | mv56_y.nii.gz | uint16 | uint8 | 512 | 512 | 11 | 0.994 | 0.994 | 5 | rat |
| 57 | mv57.nii.gz | mv57_y.nii.gz | uint16 | uint8 | 512 | 512 | 13 | 0.994 | 0.994 | 2 | rat |
| 58 | mv58.nii.gz | mv58_y.nii.gz | uint16 | uint8 | 512 | 512 | 25 | 0.994 | 0.994 | 2 | rat |
| 59 | mv59.nii.gz | mv59_y.nii.gz | uint16 | uint8 | 512 | 512 | 30 | 0.994 | 0.994 | 2 | rat |
| 60 | mv60.nii.gz | mv60_y.nii.gz | uint16 | uint8 | 512 | 512 | 56 | 0.994 | 0.994 | 2 | rat |
| index | raw | seg | raw | seg | x | y | z | x | y | z |  |
| 61 | mv61.nii.gz | mv61_y.nii.gz | uint16 | uint8 | 512 | 512 | 35 | 0.994 | 0.994 | 2 | rat |
| 62 | mv62.nii.gz | mv62_y.nii.gz | uint16 | uint8 | 512 | 512 | 58 | 0.994 | 0.994 | 5 | mouse |
| 63 | mv63.nii.gz | mv63_y.nii.gz | uint16 | uint8 | 512 | 512 | 54 | 0.994 | 0.994 | 5 | mouse |
| 64 | mv64.nii.gz | mv64_y.nii.gz | uint16 | uint8 | 512 | 512 | 30 | 0.994 | 0.994 | 5 | mouse |
| 65 | mv65.nii.gz | mv65_y.nii.gz | uint16 | uint8 | 512 | 512 | 31 | 0.994 | 0.994 | 5 | mouse |
| 66 | mv66.nii.gz | mv66_y.nii.gz | uint16 | uint8 | 512 | 512 | 10 | 0.994 | 0.994 | 5 | mouse |
| 67 | mv67.nii.gz | mv67_y.nii.gz | uint16 | uint8 | 512 | 512 | 9 | 0.994 | 0.994 | 5 | mouse |
| 68 | mv68.nii.gz | mv68_y.nii.gz | uint16 | uint8 | 512 | 512 | 106 | 0.994 | 0.994 | 2 | mouse |
| 69 | mv69.nii.gz | mv69_y.nii.gz | uint16 | uint8 | 512 | 512 | 37 | 0.994 | 0.994 | 5 | mouse |
| 70 | mv70.nii.gz | mv70_y.nii.gz | uint16 | uint8 | 512 | 512 | 15 | 0.994 | 0.994 | 5 | mouse |

### File conversions

Image volumes were converted to the NIfTI (.nii) file format to make segmentation model protoyping faster, as it is commonly used in neuroimaging and ML frameworks, such as MONAI (https://monai.io/). In MONAI, users can create dataloaders that are customized for their data formats by using Python libraries [such as `tifffile` (https://pypi.org/project/tifffile/), `python-bioformats` (https://pypi.org/project/python-bioformats/), and `pyometiff` (https://github.com/filippocastelli/pyometiff)]. In the future, we plan to develop a dataloader to allow direct use of microscopy formats, skipping the NIfTI conversion. Here, we provide the code to convert Olympus files (.oib and .oir) to NifTI (.nii) format, with metadata encoded in the NifTI1 header format. NifTI files were further compressed as .gz archive files (.nii.gz). The original Olympus files are 12-bit, and the exported NifTI files are saved as 16-bit images, as a 12-bit data type is not available. The code also provides options to export each channel separately in multichannel image volumes, separate time volumes as single volumes, and remove top and bottom slices. Further details can be found in the GitHub repository https://github.com/ctpn/minivess.

### Ground-truth annotation

#### Pre-processing

To create segmented image volumes, images were first preprocessed in Python. Single channel image volumes were individually processed using histogram equalization, Gaussian filtering, morphological operators, and thresholded into binary images. If present, image slices with poor SNR were removed from the top of a stack. Fine-tuning of binary images was achieved using 3D Slicer^45^. A general workflow of the pipeline to achieve ground-truth annotations is shown in Figure 3.

**Figure 2.**
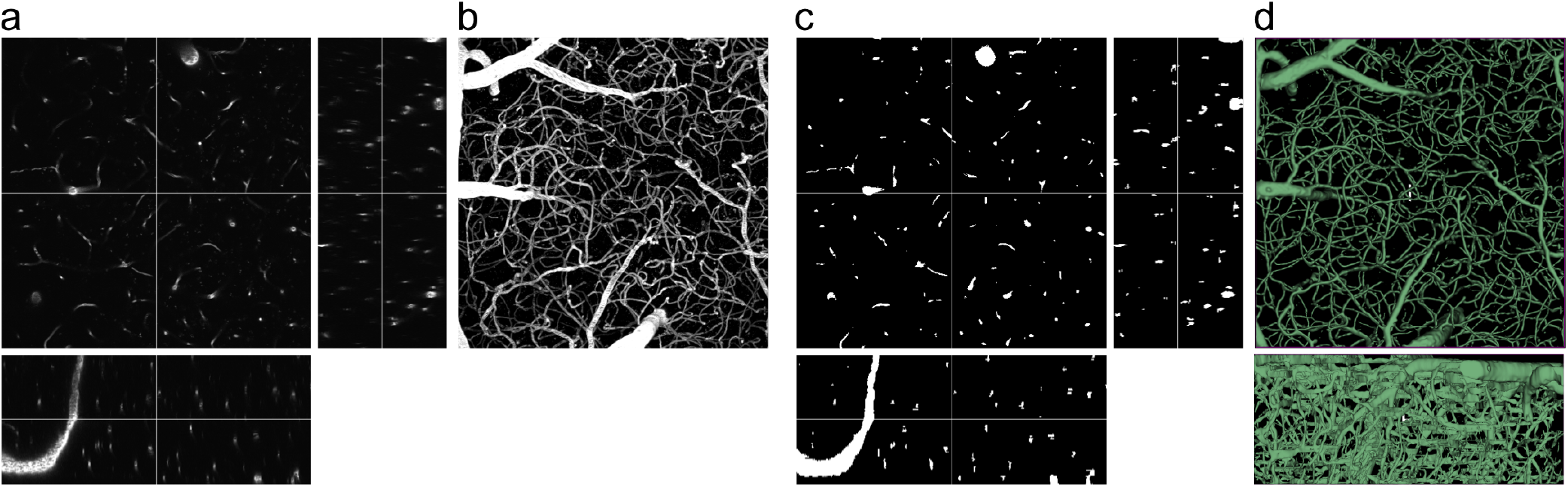
Example of a raw and segmented image volume pair in the MiniVess dataset. Orthogonal views and maximum projections of (a,b) raw and (c) segmented image volumes. (d) 3D visualization of the whole image volume in (X,Y) and (X,Z) views using 3D Slicer.

**Figure 3.**
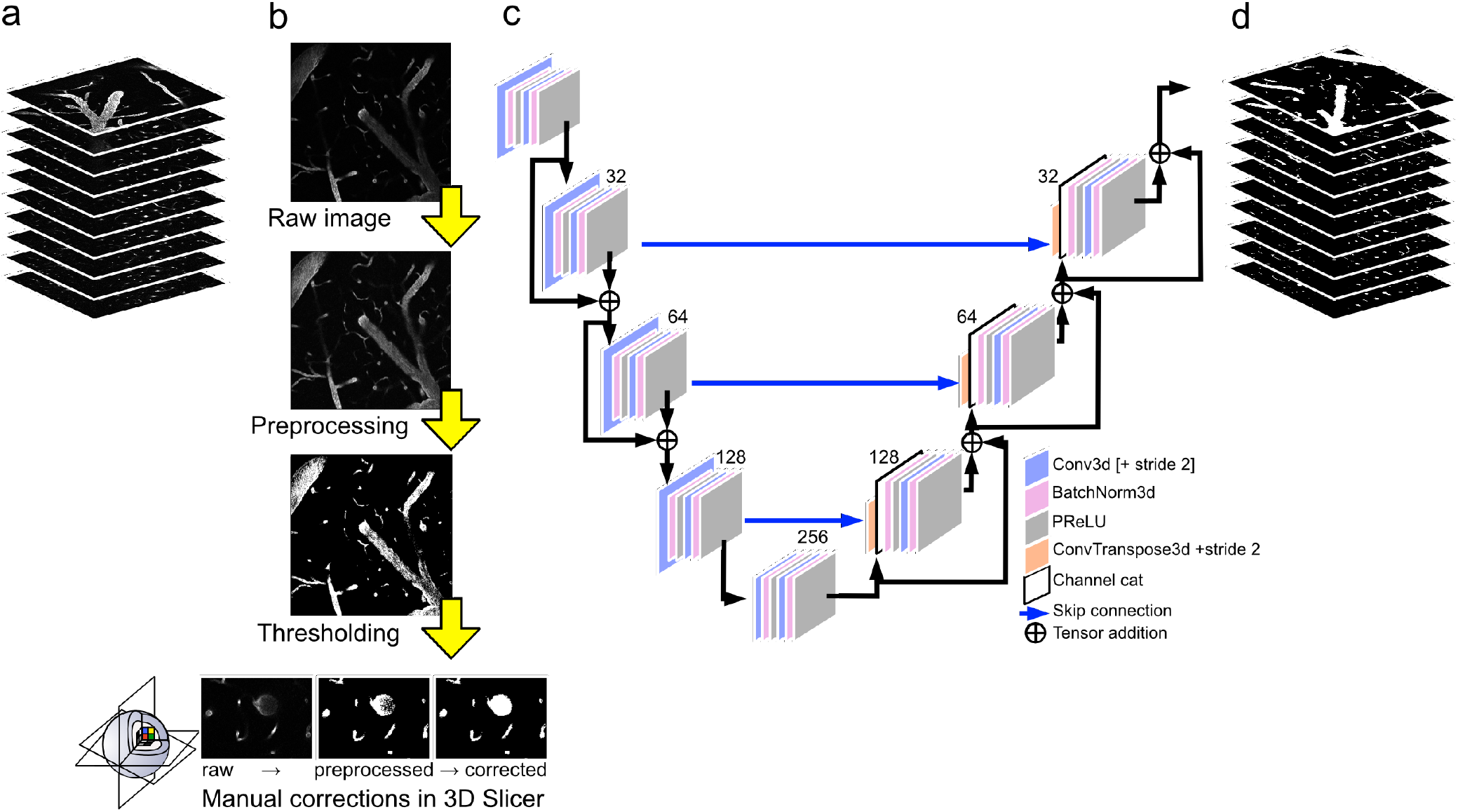
Blood vessel segmentation workflow. a) Raw image volumes from 2PFM imaging went through a series of b) preprocessing steps, followed by manual corrections conducted in 3D Slicer^45^. c) These segmented image volumes were further refined by using a 2D U-Net, which outputs d) segmented image volumes. Raw and segmented image volumes, and code for preprocessing and U-Net workflows are provided in the MiniVess project.

#### Machine learning

To improve segmentations, a 2D U-Net^23^ was trained using raw images and the preprocessed, segmented images. The U-Net consisted of 5 channels, consisting of 16, 32, 64, 128, and 256 filters, a stride of 2, batch normalization, Adam optimization (1e-4 learning rate), and the Dice loss function. Outputs from the U-Net were refined through manual corrections in 3D Slicer. Manual corrections were kept to a minimum to ensure consistency in labels within each volume. Emphasis was also placed on removing spurious noise and conserving smooth boundaries. Final segmented volumes are the result of five rounds of 2D U-Net and manual corrections in 3D Slicer (Figure 5). Supervised learning was implemented using the PyTorch-based MONAI framework^46^.

**Figure 4.**
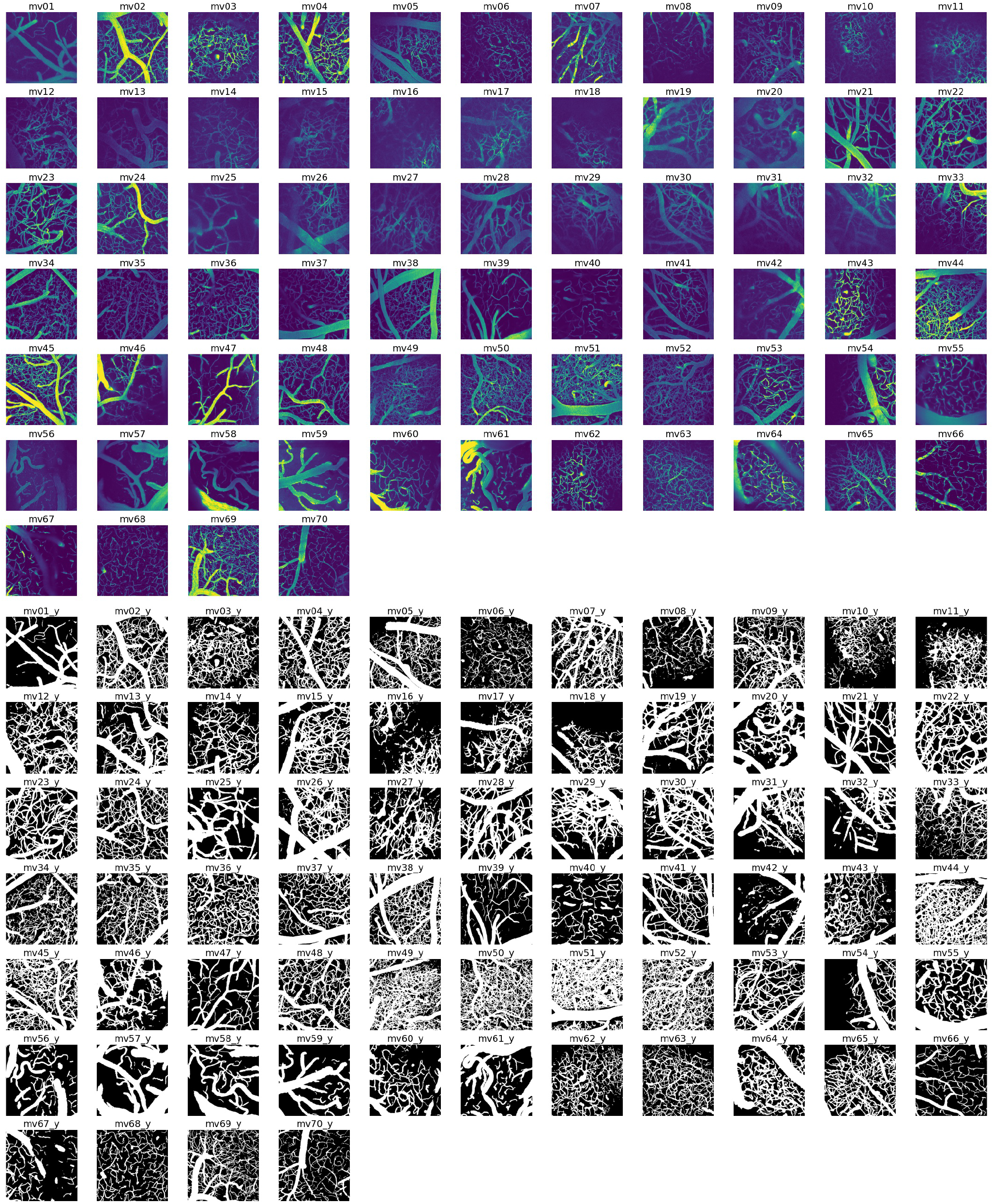
MiniVess volumes. Maximum projection images of all raw and segmented (’y’) image volumes of the MiniVess data are shown for navigation purposes. For clarity, maximum projection images consist of a maximum of 30 slices in each volume. Dark regions within the image volume that appear to have no blood vessels (e.g. diagonal in *mv16*, top of *mv18*) likely have blood vessels, but are difficult to see due to ’shadows’ cast by larger blood vessels above, which are not included in the image volume.

**Figure 5.**
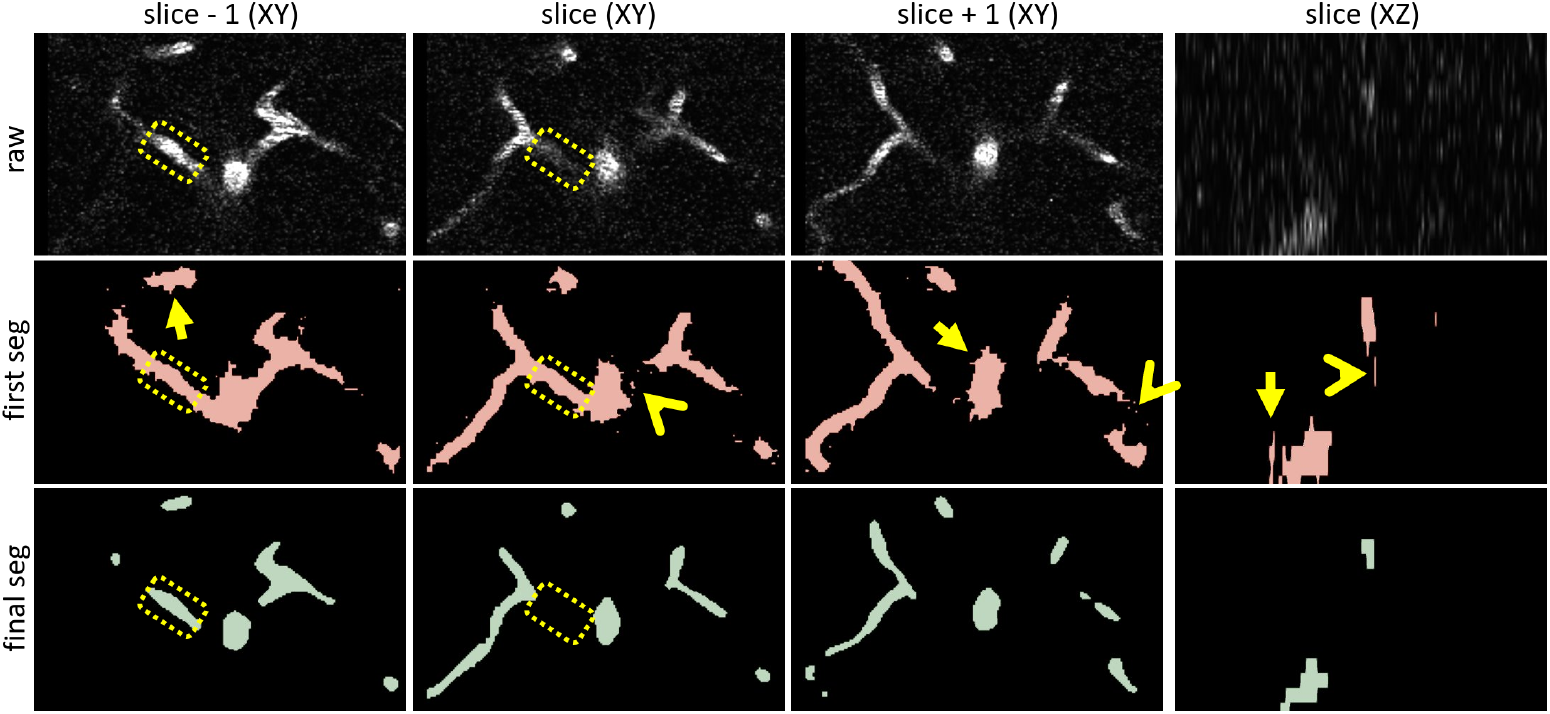
Validation of segmentations. Cropped images from the XY and XZ planes taken from the raw volume (’raw’), and the first (’first seg’) and final (’final seg’) rounds of segmentations, are shown for comparison (image volume *mv18*). In addition to the slice-of-interest (’slice (XY)’ and ’slice (XZ)’), the image slices above (’slice - 1 (XY)’) and below (’slice + 1 (XY)’) are also shown. In the first segmentation, vessel edges are less uniform (arrows), spurious noise is evident in the segmentations (angles), and vessel segments that are outside of the slice-of-interest, but present in the slices above or below, are included in the segmentation of the current slice (compare dotted outlines). In contrast, in the final segmentation, vessel edges are more uniform, and segmentations are closer to what is visible in the current slice only, according to acquisition parameters. Each round of segmentation consisted of manual corrections and U-Net outputs. For manual corrections of U-Net outputs, emphasis was placed on minimizing manual drawing to reduce human error, and smoothing edges. For example, jagged borders (arrows) observed in the first round of segmentation are smooth by the final segmentation.

### Data Records

The data is currently stored in the (EBRAINS) repository in compressed NiFTi format (*.nii.gz). Each raw image stack has an annotated equivalent, designated by a ’y’ in the file name. Details for each image can be found in the metadata, encoded in the NIfTI1 header format. Each image stack represents a different field-of-view in the cerebrovasculature. Information specific to each image stack can be found in Online-only Table 1. Maximum projection images of all image volumes are shown in Figure 4.

### Technical Validation

Image volumes were collected and curated by CP (7 years of experience). Ground truth annotations were achieved by using classic image processing tools (see Methods), manual annotations by CP, and a 2D U-Net^23^. Accuracy of the final annotations were qualitatively confirmed by CP, and then independently confirmed by MFR and HS (Figure 2). Final segmentations are the result of 5 rounds of manual annotations or corrections and outputs of the U-Net. A comparison between rounds of segmentations can be found in Figure 5.

### Usage Notes

MiniVess contains image volumes of cerebrovasculature from wild-type mouse, transgenic mouse, and transgenic rat brains. Although small in size, the variety of background strains and species in the MiniVess dataset represents rodent strains that are commonly used in wet labs.

The dataset can be downloaded as NiFTi (.nii.gz) files which can then be easily uploaded into machine learning models, or manipulated using FIJI (Fiji Is Just ImageJ), Python, MATLAB, etc. We provide a tutorial of how to use the MiniVess dataset in a U-Net, built in the MONAI framework (https://github.com/project-monai/monai).The MONAI framework also provides several tutorials using NiFTi images, which can be further explored using the MiniVess dataset.

By making the raw and annotated data available, we hope that the MiniVess dataset can be used as a validation dataset by those evaluating their supervised, semi-supervised, or unsupervised segmentation models, and assist the field to use more data-centric ways to design and evaluate their segmentation models.

## Code availability

We provide the Python code to separate multichannel and time series 2PFM image volumes into single volumes, which are easier to manipulate. multichannel XY, XYZ, XYT, and XYZT images are supported. For multichannel images, the user will be asked to select the channel of interest to export. For images with multi-T volumes (XYT and XYZT), the user has the option of exporting each T-stack separately, or as a single file. The code can be accessed at our Github repository https://github.com/ctpn/minivess.

## Acknowledgements

This work was supported by funding from the National Institute of Biomedical Imaging and Bioengineering of the National Institutes of Health (RO1-EB003268, awarded to K.H.), the Canadian Institutes of Health Research (FDN 154272, awarded to K.H.), and the Temerty Chair in Focused Ultrasound Research at Sunnybrook Health Sciences Centre. This work was also supported by the program for Brain Mapping by Integrated Neurotechnologies for Disease Studies (Brain/MINDS) from the Japan Agency for Medical Research and Development AMED (JP15dm0207001).

## Author contributions statement

PT conceived the experiment. CP and PT wrote code for data conversion and the U-Net. CP conducted the experiments and analyzed the results. CP and PT wrote the manuscript. All authors reviewed the manuscript.

